# Axolotl epigenetic clocks offer insights into the nature of negligible senescence

**DOI:** 10.1101/2024.09.09.611397

**Authors:** Yuliia Haluza, Joseph A. Zoller, Ake T. Lu, Hannah E. Walters, Martina Lachnit, Robert Lowe, Amin Haghani, Robert T. Brooke, Naomi Park, Maximina H. Yun, Steve Horvath

## Abstract

Renowned for their regenerative abilities, axolotls also exhibit exceptional longevity, resistance to age-related diseases and apparent lack of physiological declines through lifespan, and have thus been considered organisms of negligible senescence. Whether axolotls display epigenetic hallmarks of ageing remains unknown. Here, we probe the axolotl DNA methylome throughout lifespan and present its first epigenetic clocks. Both at tissue-specific or pan-tissue levels, the clocks are biphasic, capable of predicting age during early life but not for the rest of its lifespan. We show that axolotls exhibit evolutionarily conserved features of epigenetic ageing during early life, yet their methylome is remarkably stable across lifespan, including at Polycomb Repressive Complex 2 (PRC2) target sites, suggesting that this species deviates from known patterns of epigenetic ageing. This study provides molecular insights into negligible senescence and furthers our understanding of ageing dynamics in animals capable of extreme regeneration.

## Introduction

Salamanders such as the axolotl (*Ambystoma mexicanum*) are the evolutionarily closest organisms to humans capable of regenerating extensive sections of their body plan, including parts of their eyes, lungs, heart, brain, spinal cord, tail and limbs throughout their lives^1–3^, constituting valuable models for regeneration studies. Yet, urodele amphibians are also characterised by an apparent lack of physiological declines through lifespan^4,5^, indefinite regenerative capacity^6^, extraordinary longevity^5,7^, and defiance of the Gompertz law of mortality^5,8,9^, key features of negligible senescence. Their long lifespans and lack of experimental tractability have historically restricted their use in ageing studies. However, recent technological advances have enabled the axolotl as a tractable model system, with a toolkit including transgenesis and genome editing^10,11^, genome assemblies^12–14^, transfection^15,16^ and imaging^17^ techniques, and tools to probe age-related processes including cellular senescence^18,19^ and telomere biology^20^. This, combined with its relatively short lifespan within Urodela (average lifespan c. 10-13 years, with sexual maturation at 10-12 months), has transformed the axolotl into an emerging model for ageing research^5^.

Throughout life, axolotls exhibit several age-defying traits, including dermal thickening^21^, progressive skeletal ossification^21,22^, and cancer resistance^23^. Further, their tissues do not accumulate senescent cells with age^5,24^, thereby circumventing a major hallmark of ageing and driver of age-related disorders^25^, in keeping with their proposed negligible senescence status. Yet, whether axolotls exhibit signs of molecular ageing remains unknown.

Changes in the methylation level of cytosines (5-methylcytosine) within CpG dinucleotides constitute a primary hallmark of molecular ageing^26^. Indeed, age-related changes in DNA methylation (DNAm) occur across animal species, including mammals^27,28^, birds^29,30^, fishes^31,32^ and amphibians^33–35^. More recently, the Mammalian Methylation Consortium has confirmed that age-related gains in methylation can be observed at target sites of the Polycomb Repressive Complex 2 (PRC2), which catalyses the tri-methylation of lysine 27 on histone H3 (H3K27me3) in all mammalian species^36^. DNA methylation at PRC2 target sites may constitute a universal biomarker of aging and rejuvenation in mammalian systems^37^.

Multivariate regression models based on the methylation status of multiple CpG sites provide accurate age estimators, commonly known as ‘epigenetic clocks,’ in mammalian species^38–41^. Initially restricted to humans and mice, the identification of highly conserved CpGs facilitated the construction of multispecies clocks such as the pan-mammalian clock, capable of age predictions across all mammalian species^36^. This included exceptionally long-lived (e.g. bats^42^) and slow ageing (e.g. naked mole rat^43^) mammals. More recently, this has extended to African clawed frog species-*Xenopus laevis* and *Xenopus tropicalis*-^33,34^. Of note, the generation of these clocks was achieved through an Infinium methylation array platform based on highly conserved mammalian CpGs^33,34,44^, supporting the evolutionary conservation of age-related epigenetic signatures between amphibians and mammals, and demonstrating the applicability of mammalian methylation arrays in amphibians.

Here, we conduct DNA methylation profiling of axolotl tissues at CpGs associated with ageing across mammalian and amphibian species. We develop axolotl epigenetic clocks at both pan- and single tissue levels and uncover that axolotls exhibit conserved epigenetic ageing traits during early life but not thereafter, deviating from the established notion of organismal ageing. We reveal that, in contrast to mammals, the axolotl methylome is remarkably stable and does not exhibit substantial shifts at either global or PRC2-associated gene levels late in life. These findings offer critical insights into the nature of negligible senescence.

## Results

### Axolotl epigenetic clocks are biphasic and can only predict age during early life

To construct axolotl epigenetic clocks, we generated methylation profiles for 180 axolotl samples (Supplementary Table 1) from individuals ranging from 0.08 (4 weeks) to 21 years of age, encompassing 6 tissue types (limb, tail, skin, liver, spleen, and blood). Profiling was conducted using the HorvathMammalMethylChip40 Infinium array platform, which informs on the methylation status of highly-conserved CpGs used to track age across mammals^27,36,44^ and was previously employed to build accurate age predictors for amphibians, namely the anurans *Xenopus tropicalis* and *Xenopus laevis*^33,34^. Only conserved CpGs that mapped to the axolotl genome (AmexG_v6.0-DD assembly, n=5386 CpGs) were used in our study.

Unsupervised hierarchical clustering of all axolotl samples (Extended Data Fig. 1) did not reveal clustering based on tissue type or sex, suggesting that interindividual effects have a more pronounced influence, in line with prior reports on DNA methylation patterns in amphibians and reptiles^45^. In contrast, mammalian methylation studies show that tissues have very distinct methylation patterns, resulting in distinct clusters^27^.

Next, we leveraged our datasets to build DNAm-based age estimators using elastic net regression models (in which chronological age acts as dependent variable and CpG methylation levels as covariates). To obtain unbiased estimates of the DNAm clocks, we performed leave-one-out (LOO) cross-validation of the training dataset, arriving at estimates of the age correlation R-Pearson correlation between the estimated DNA methylation age (DNAm age) and the chronological age- and the median absolute error, MAE, based on the absolute difference between predicted and actual age in units of years. Surprisingly, all attempts to build an axolotl DNAm clock using datasets spanning the entire age range were unsuccessful (Fig. 1a-d), as indicated by the poor correlation coefficients obtained for either single tissue or pan tissue clocks.

**Figure 1.**
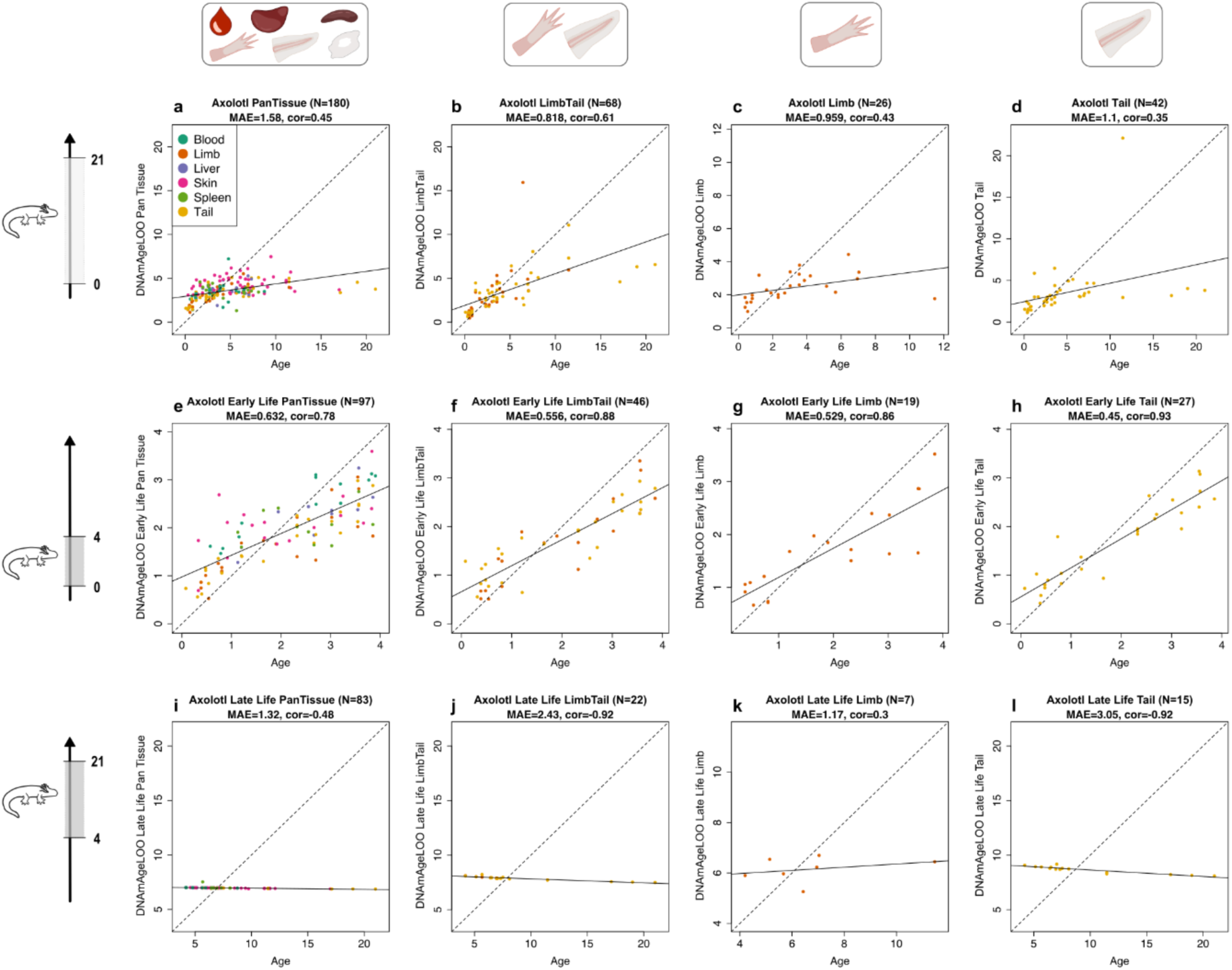
Axolotl epigenetic clocks are biphasic and can only predict age during early life. Cross-validation study of epigenetic clocks for *Ambystoma mexicanum*. Leave-one-sample-out (LOO) estimate of DNA methylation age (y-axis, in units of years) versus chronological age (x-axis, units in years) for *A. mexicanum* samples across selected range of age. Dots are coloured by tissue type. **a**-**d**, Epigenetic clocks across all ages (up to maximum 21 years): **a**, pan-tissue, **b**, limb-tail, **c**, limb, **d**, tail. **e**-**h**, Epigenetic clocks across early life cohort (<4 years): **e**, pan-tissue, **f**, limb-tail, **g**, limb, **h**, tail. **i**-**l**, Epigenetic clocks across late life cohort (>4 years): **i**, pan-tissue, **j**, limb-tail, **k**, limb, **l**, tail. Each panel reports the total sample size (N), median absolute error (MAE), in years, Pearson correlation coefficient (cor).

We then asked whether it would be possible to build a clock for a particular period of the axolotl lifespan. Indeed, we were able to build accurate DNAm age estimators for samples corresponding to axolotls up to 4 years of age (a life period hereby designated as ‘early life’, Fig. 1e-h and Extended Data Fig. 2). Pan tissue, limb, tail, and the combination of limb and tail clocks produced strong Pearson correlation coefficients (R near or above 0.8, Fig. 1e-h), with single tissue clocks from limb (MAE=0.529) and tail (MAE=0.45) being more accurate (Fig. 1f-h). This was also the case for blood and liver, while smaller coefficients were observed for skin and spleen (Extended Data Fig. 2). To confirm that the clocks can respond to induced changes in global methylation levels, we exposed axolotl cells to the demethylation agent 5-aza-2’-deoxycytidine, known as decitabine (DAC). As expected, increasing doses of DAC led to concomitant declines in overall methylation levels as estimated by either ELISA-based global methylation assay (Extended Data Fig. 3a) or by the Infinium methylation arrays (Extended Data Fig. 3b). This resulted in changes in DNAm age as estimated by the corresponding clock (Extended Data 3c). Further, we confirmed that the early life clocks were able to predict age of independently generated limb and tail samples with reasonable accuracy (MAE=0,172 (Pan Tissue), 0.255 (Limb Tail), Extended Data 3d-e), validating their use as axolotl age predictors. Strikingly, it was not possible to build DNAm clocks based on samples from individuals over 4 years of age, despite analysing over 80 samples of good quality and tissue representation, at neither single nor pan-tissue levels (Fig. 1i-l). Collectively, these data indicate that axolotl epigenetic clocks exhibit a biphasic nature: they can predict age during early life, yet they stop ticking for the rest of the axolotl’s lifespan.

### Dual-species DNAm clocks during axolotl early life

To explore the presence of conserved age-related epigenetic signatures between axolotls and frogs, we first investigated whether it was feasible to construct dual-species epigenetic clocks. We hypothesized that if a subset of CpG sites showed consistent age-related methylation patterns in both amphibian species, it would be possible to create a combined species predictor. Indeed, we successfully developed dual-species axolotl-frog clocks by integrating our axolotl methylation profiles—spanning multiple tissues or limb and tail samples up to 4 years old—with a published pan-tissue frog dataset^33^ generated using the same methylation array (Fig. 2a-c). These results show that the age-related CpG methylation changes observed during the early life stages of axolotls are at least partially conserved across amphibians.

**Fig. 2.**
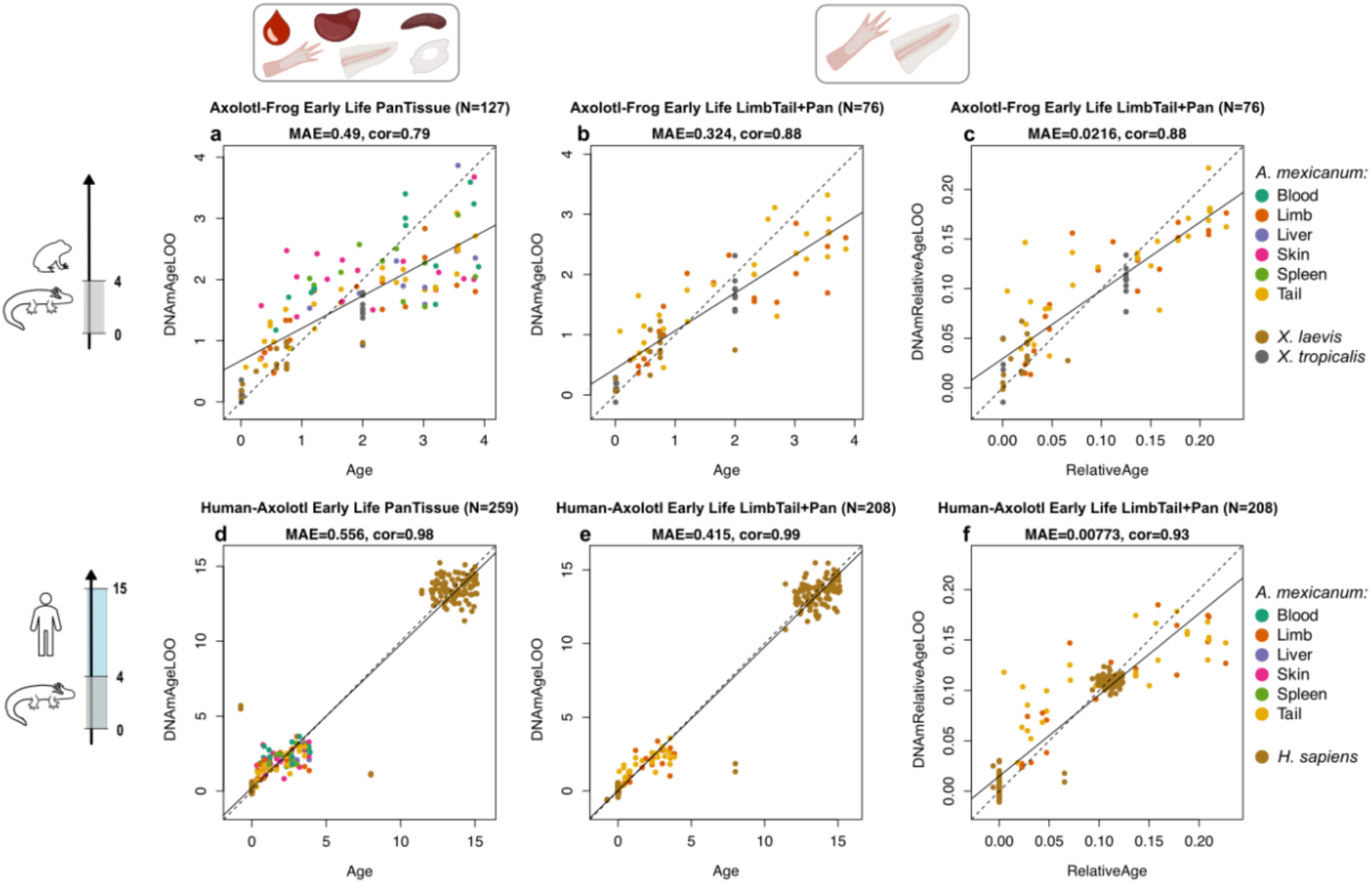
Dual species Axolotl-*Xenopus* and Axolotl-Human epigenetic clocks reconstitute a biphasic behaviour and predict age during early life. Cross-validation study of epigenetic clocks for *Ambystoma mexicanum*, *Xenopus laevis*, *Xenopus tropicalis*, and *Homo sapiens*. Leave-one-sample-out (LOO) estimate of DNA methylation age (y-axis, in units of years) versus chronological age (x-axis, units in years) for samples across selected range of age. Dots are coloured by species and tissues. **a**-**c**, Epigenetic clocks (<4 years for *Ambystoma mexicanum* [PanTissue N = 97, LimbTail N = 46], *Xenopus laevis* and *tropicalis* [PanTissue N = 30]) for **a**-**b**, chronological age, and **c**, relative age. **d**-**f**, Epigenetic clocks (<4 years for *Ambystoma mexicanum*, <15 years for *Homo sapiens* [PanTissue N = 162]) for **d**-**e**, chronological age, and **f**, relative age. Each panel reports the total sample size (N), median absolute error (MAE), in years, Pearson correlation coefficient (cor).

We then asked if conserved trends could also be found between axolotl and mammals. In support of this notion, we were able to build accurate dual species axolotl-human clocks, where the aforementioned axolotl datasets together with available human methylation profiles^46^ were used for training (Fig. 2d-f). In addition to developing clocks that estimate chronological age, we also built relative age predictors to account for differences in species lifespan (Fig. 2c and 2f). Here relative age was defined as ratio between chronological age and maximum lifespan. Comparing the predicted relative age on the common scale, we noticed a trend of axolotl age overestimation (until relative age 0.1) in relation to both *Xenopus* species and humans. This may correspond to the temporal acceleration of methylation changes during early life, appearing as a species-specific pattern. Of note, we were unable to build dual clocks with either species when using the axolotl methylation data from individuals over 4 years of age, despite the fact that clocks spanning the entire lifespan can be constructed for frogs and humans combined with other species^33,36,43^. Together, these findings have several implications. First, they suggest that axolotls until 4 years of age undergo age-related epigenetic changes that are shared, at least in part, with other amphibians and even mammals, which exhibit organismal ageing. Second, they suggest that, for the rest of their lifespan, axolotls display divergent epigenetic patters and thus deviate from the conventional notion of epigenetic ageing.

### Features of age-related CpGs during axolotl early life

To uncover the nature of the CpGs whose methylation changes are most highly correlated with chronological age in axolotls, we performed an epigenome-wide association study (EWAS) of age for tissue samples during the early life period (Fig. 3a-g). Using a genome-wide significance cut off of p<1.0E-04 we identified a number of CpGs, whose amount and identity varied across tissues. The majority of significant CpGs were captured in limb and tail tissues (Fig. 3a-b), followed by blood (Extended Data Fig. 4). No significant CpG changes were detected in liver, skin or spleen at our adopted threshold (Extended Data Fig. 4). For the top 100 age-related CpGs, demethylation events are somewhat prevalent in each tissue except for skin, whose associated CpGs exhibit a trend towards gaining methylation with age (Fig. 3a-b; Extended Data Fig. 4).

**Fig 3.**
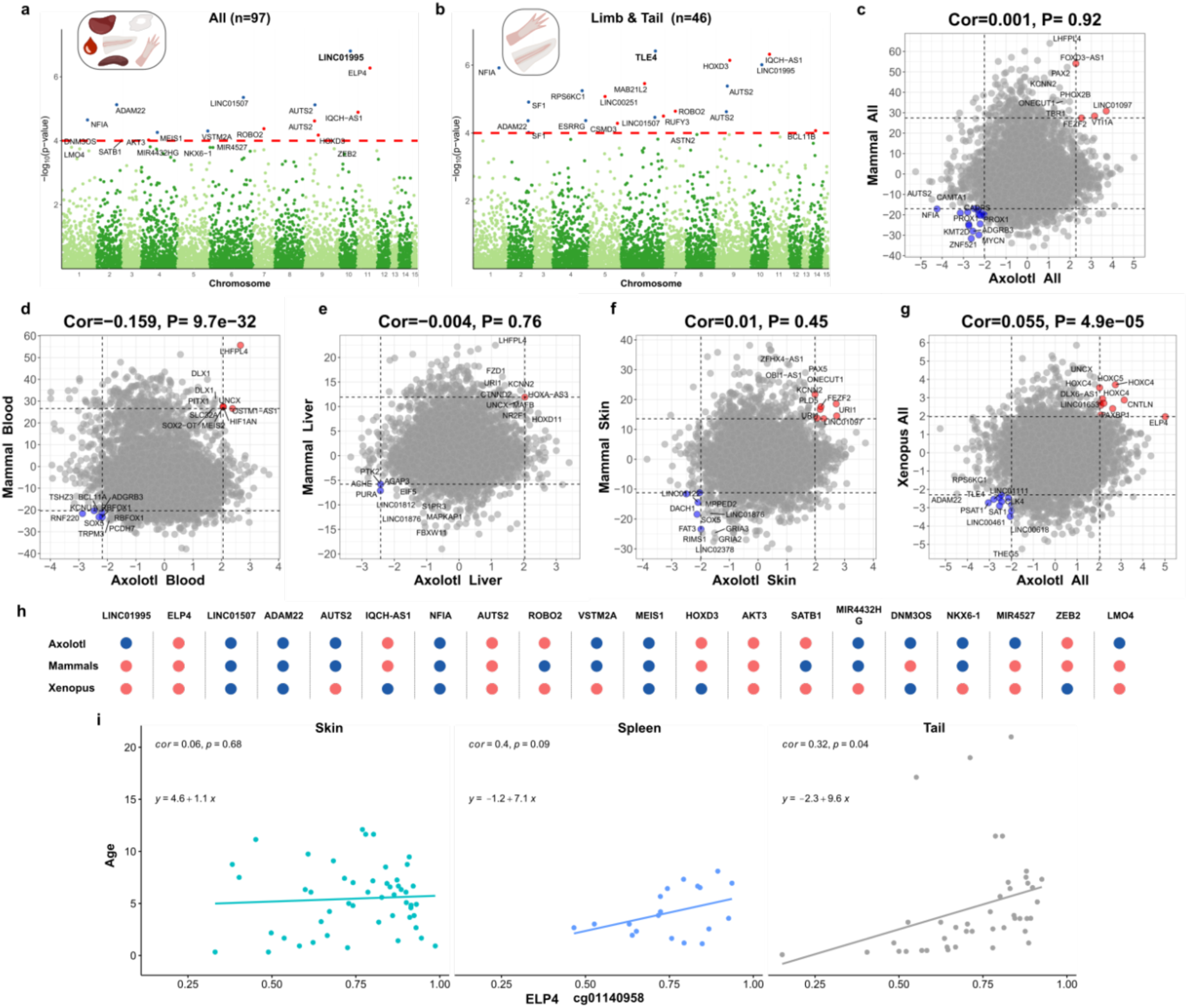
Meta-analysis of age-related methylation change during axolotl early life reveals alterations related to mammalian and amphibian ageing. **a**-**b**, Manhattan plots of the epigenome-wide association (EWAS) of chronological age in *A. mexicanum* tissues across early life adult group (<4 years): **a**, pan-tissue, **b**, limb-tail. The red dash lines indicate suggestive levels of significance at P < 1.0E-04. CpGs in red and blue reflect positive and negative correlation with age, respectively. The y-axis displays -log base 10 transformed P-value and the x-axis displays chromosome number based on the *A. mexicanum* genome (v. 6.0-DD). Labels are provided for the top 20 hypermethylated / hypomethylated CpGs according to the product of Z scores in x- and y-axis. Each panel reports the total sample size (n). **c**-**f**, Axolotl versus Eutherian EWAS of age and, **g**, axolotl versus *Xenopus* EWAS of age, based on Z statistics. Dots correspond to CpGs that are present on mammalian array and map to the *A. mexicanum* genome. Each panel reports Pearson correlation and its P-value between the two EWAS Z statistics. Labels are provided for the top 10 hypermethylated/hypomethylated CpGs according to the product of Z scores in x- and y-axis. **h**, The methylation state of the top 20 age-related CpGs in axolotls, in comparison with mammals and *Xenopus*. Red and blue colours denote hypermethylated and hypomethylated CpGs, respectively. **i**, The distribution of individual CpG methylation across ageing in *A. mexicanum* skin, spleen and tail. The panel reports Pearson correlation coefficient (cor) and P-value (p).

Interestingly, the most significant age-correlated CpGs (Fig. 3a-b; Extended Data Fig. 4; Supplementary Table 3) were associated to gene loci that include developmental transcription factors of the homeobox category or their regulators, as well as genes previously linked to ageing. At pan tissue, limb and tail levels these include the homeobox regulator and Polycomb binding factor *AUTS2*^47^, linked to skin ageing^48^, the homeobox transcription factors *HOXD3* and *MEIS1*, the transcriptional co-repressor *TLE4*, the long non-conding RNA *LINC01995*, a putative *SOX2* modulator^49^, and developmental regulators such as the metalloprotease *ADAM22*. For blood, associated genes include the transcription factors *PAX2* and *MAFB*, and the age-related hypertension factor *NEDD4L*^50^.

A comparison between the results from our axolotl EWAS and equivalent data derived from eutherian mammals^35^(Mammalian Methylation Consortium^35^) or *Xenopus*^33^ suggests a low correlation overall (Fig. 3c-g), with changes in top age-related genes closer to frog (Cor = 0.055) than to mammals (Cor = 0.001). However, it also highlights shared methylation trends for key CpGs, such as those associated with the developmental genes *HOXC4, HOXC5* and *UNCX,* which gain methylation with age in both axolotls and frogs (Fig. 3g). Interestingly, *UNCX*, top gene with positive CpG methylation-age correlation in frogs^33^, also exhibits a similar methylation trend in human and axolotl blood and liver (Fig. 3d-e). Indeed, we identified several gene loci whose associated CpGs exhibit age-related methylation patterns conserved between axolotls up to 4 years of age, mammals and *Xenopus* (Fig. 3h). We also note that methylation changes are highly tissue specific, and even strongly age-associated loci do not necessarily exhibit methylation changes of equivalent magnitude in different tissues. As an example, the methylation level of a conserved-axolotl, frogs, mammals-CpG site associated to *ELP4,* exhibits noticeable variations across tissues (Fig. 3f). Together, these observations support the existence of conserved features of age-related methylation between mammals, frogs, and axolotls in their early life period.

To further explore the biological features of axolotl age-associated CpGs, we conducted GREAT functional enrichment analysis^51^ based on the top 150 positively and negatively age-correlated CpGs (Fig. 4). While few enrichments were observed for negatively age-correlated CpGs, significant enrichments were found for those with positive correlations. Among these, a large proportion falls into the category of developmental and morphogenetic processes (Fig. 4) and include homeobox/homeodomain transcription factor gene categories, consistent with our EWAS study. For example, positively correlated CpGs are enriched in *HOXL* regions, resonating with recent findings which report that enrichment in methylated CpGs within *HOXL* is a feature of long-lived species^27^. This is additionally supported by gene ontology (GO) analysis, in which terms related to developmental processes and *HOX* genes in particular feature prominently alongside transcriptional processes (Extended Data Fig. 5a). These observations are in agreement with prior findings in *Xenopus*^33^ and mammals^36^, reinforcing the notion that until 4 years of age, axolotls display epigenetic ageing trends similar to those found in species with organismal senescence.

**Figure 4.**
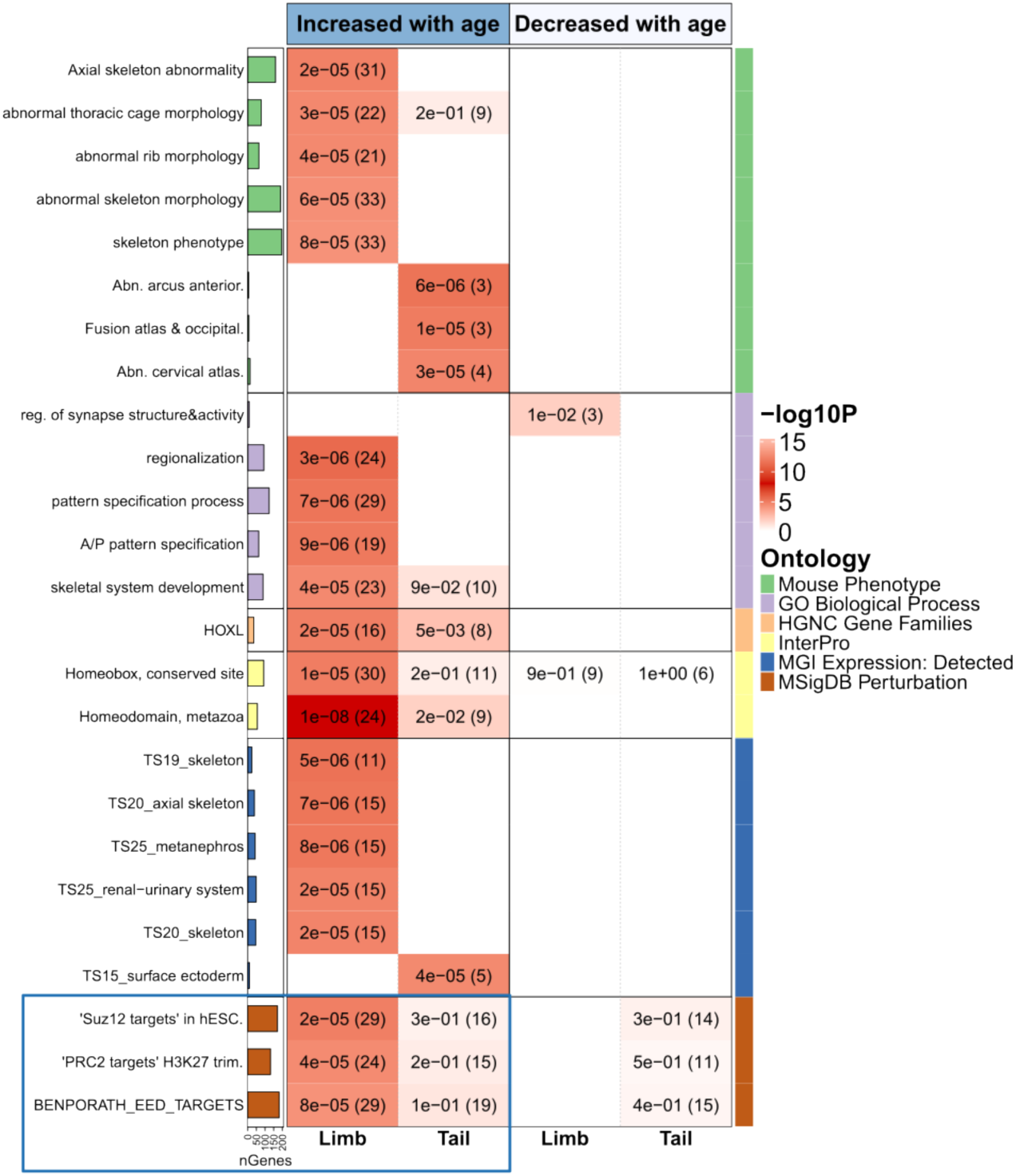
Functional enrichment of EWAS meta-analysis. GREAT functional enrichment results for top 150 age-related CpGs from EWAS in 1) limb, 2) tail. The y-axis denotes the name of a functional gene set/biological pathway. Each panel represent the enrichment results based on the top 150 CpGs with positive or negative age correlation. The heatmap colour gradient is based on -log10 (unadjusted hypergeometric P-value) multiplied by the sign of OR > 1. Abbreviations: Abn. – abnormal, TS - Theiler stages of embryonic development, BENPORATH_EED_TARGETS denotes EED targets - target genes of the Polycomb protein EED (GeneID=8726), Suz12 targets – target genes of the Polycomb protein SUZ12 (GeneID=23512) identified by ChIPSeq in human embryonic stem cells.

The analysis also highlighted points of departure from other organisms which could pertain to axolotl biology. These include an enrichment in abnormal skeleton morphology and related terms, which may be explained by intrinsic differences in skeleton development and continuous ossification^22^. Further, we note a distinct lack of enrichment in mortality, disease and cancer categories, as opposed to other species^33,36^. The latter is consistent with prior reports of cancer resistance in axolotls^23^, and defiance of Gompertz law of mortality among salamander species^5,9^. While the absence of enrichment could also be explained by an exclusion of the associated CpGs due to lack of alignment to the axolotl genome, we note that categories related to cancer and CNS abnormalities feature prominently among age-related CpGs in frogs, which exhibit a reduced set of mapped CpGs compared to axolotls^33^.

Interestingly, we also observed a slight enrichment in the targets of the PRC2 component Embryonic Ectoderm Development (EED) for limb tissues, and tail to a lesser extent (Fig. 4). CpGs related to PRC2 targets have been widely reported to gain methylation with age across species, from mammals^36,37,42,52,53^ to amphibians^33,34^, constituting a hallmark of epigenetic alterations with age. Yet, the enrichment significance for axolotl CpGs is comparatively low than for other species^33,36^. This is further supported by human chromatin state analysis of age-related CpGs (Extended Data Fig. 5b), which indicates limited overlap between positively age-correlated CpGs and PRC regions and bivalent promoters with the exception of limb tissues. This analysis is based on human data, which invites caution. Nevertheless, these observations are noteworthy given the differences between axolotl and other amphibian species. This is exemplified by skin, for which positively age-related CpGs are heavily enriched in PRC2 target sites in *Xenopus*^33,34^, yet no enrichment -and even the opposite trend-is observed in axolotls (Extended Data Fig. 5b). This may represent an important evolutionary divergence, of relevance to the negligible senescence traits exhibited by axolotls.

### Axolotls exhibit stable global and PRC2 targets-associated DNA methylation throughout lifespan

The absence of a clock signal in axolotls beyond their early life period prompted us to investigate DNA methylation changes over larger scales. Across species, the ageing process is accompanied by local gain in DNA methylation (especially in CpG islands and PRC2 target regions) and, in certain instances, whole-genome hypomethylation^37,54–56^. To address whether this is also the case in axolotls, we measured relative changes in global methylation through ELISA, as previously employed in this species^57^. We chose the limb as representative tissue, for which we had significant amount of data on age-related processes, and selected 4 age points to cover critical stages across axolotl lifespan, from maturation to old age: 0.73 (maturing, actively growing animal), 1.1 (sexually mature), 3.55 (mature animal close to point where arrays stop tracking age-associated changes in methylation) and 9.84 (old animal, near the average lifespan for this species) years of age (Fig. 5). Relative methylation profiling revealed a lack of significant changes in global methylation across ages (Fig. 5a). While a trend towards hypomethylation is seen in individuals from 0.73 to 3.55 years of age, there is little difference between 3.55- and 9.84-year-old individuals. This suggests that axolotls, compared to other species, maintain a more stable epigenome through lifespan.

**Figure 5.**
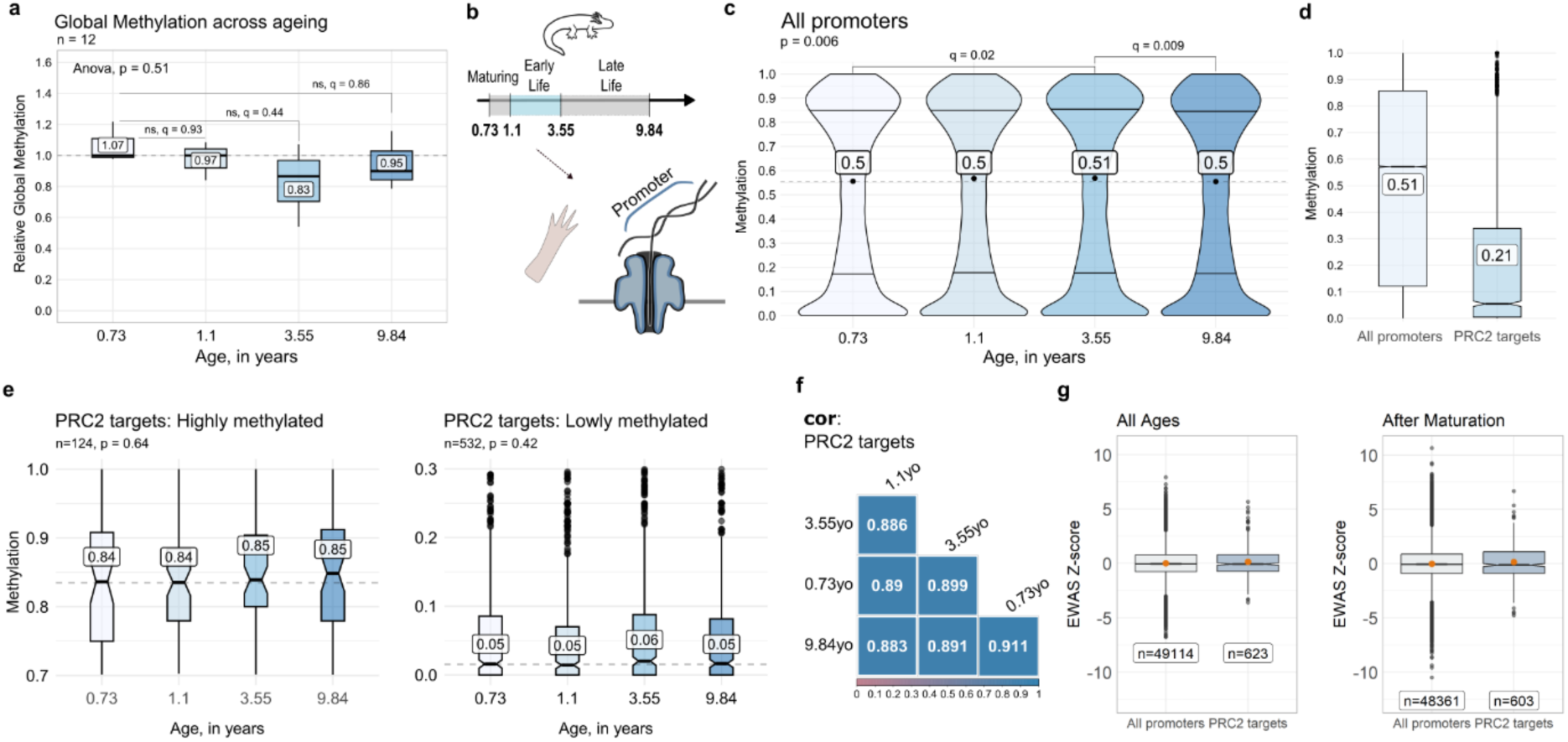
Global DNA and PRC2 target gene-specific methylation exhibit no significant changes throughout axolotl lifespan. **a**, Global methylation changes across axolotl ageing in limb. The relative global methylation level changes are indicated. The panel reports the total samples size (n), with three animals per age group. Statistical significance was evaluated by one-way ANOVA with adjustments for comparisons between all age groups and the youngest group using the Tukey method, p-value and q-values are indicated. **b**, schematic representation of the experimental set up. Promoter-targeted nanopore sequencing of axolotl limb samples with methylation calling. The age of limb samples is indicated in years. **c**-**e**, Methylation distribution in detected promoter regions across axolotl limb samples: **c**, for all promoters across age; **d**, for all promoters and Polycomb repressive complex 2 (PRC2) target genes; **e**, for highly and lowly methylated PRC2 target genes across age. PRC2 targets were defined based on human ChipSeq dataset^60,75^. Panels report gene regions number (n). Statistical significance was evaluated by Kruskal-Wallis test with adjustments for multiple comparisons using the Bonferroni method, p-values and q-values are indicated. **f**, Age pairwise Pearson correlation analysis of PRC2 target gene methylation. Colours indicate the degree of pairwise correlation. **g**, EWAS of age across all promoters and PRC2 targets. The analysis is based on all ages and on samples after maturation, excluding the youngest 0.73yo. Panels report the gene regions number (n). Box plots in **a, d, e** and **g** centre line and dots inside violin plots in **c** report median. Means are presented in white boxes. White boxes in **g** report the total sample size (n).

To further test the stability of the axolotl methylome through lifespan and achieve a more comprehensive, unbiased profiling of CpG methylation at genome scales, we employed Oxford Nanopore Technologies (ONT) for long-range 5mC methylation analysis (Fig. 5b). Due to the large size of the axolotl genome (30Gb^12^) we took advantage of adaptive sampling, which allows targeted sequencing of predetermined regions, to probe annotated promoters genome-wide. We covered samples for the same life stages used in the global methylation analysis to ensure consistency and provide independent assessment of our clock findings. As our epigenetic predictor ceases to track methylation changes after 4 years, we hypothesised the methylome to stabilise, with no significant deviations occurring between the 3.55- and 9.84-year-old samples.

Interestingly, methylation profiling across promoter regions did not reveal compelling evidence of changes in global promoter hyper or hypomethylation (Fig. 5c). This suggests that the axolotl methylome does not exhibit the common epigenetic shifts that characterise aging in other species^35,37,54,55^. We then asked whether highly (> 0.7) or lowly (< 0.3) methylated genes are more susceptible to age-related changes. By narrowing down the analysis, we did not observe a clear skewing towards hypermethylation or hypomethylation. Lowly methylated genes remained more stable, while highly methylated genes showed a slight shift towards further hypermethylation in the 3.55-year-old sample (Extended Data Fig. 6a). This contrasts with findings in mammals, in which methylation of lowly methylated genes increases significantly with age, with PRC2 target genes being particularly prone to this gain^36,37^. We thus focused on PRC2 target genes. We employed the PRC2 binding site annotation that was previously used by the Mammalian Methylation Consortium^58–60^, taking into account the high conservation of gene regulatory machinery among vertebrates^33,61,62^. We observed that PRC2 target regions display a low DNA methylation level compared to median methylation across other promoter regions (Fig. 5d), as expected^37,63,64^. Importantly, we could not detect global hypermethylation events at these regions in lowly (< 0.3) methylated genes (Fig. 5e) across ageing, but a slight gain in methylation in highly (> 0.7) methylated ones. Furthermore, we observed a strong correlation (0.88 and higher) among the samples (Fig. 5f), similar to what is seen in the comparison across all promoter regions (Extended Data Fig. 6b). Interestingly, the youngest and oldest samples showed the lowest correlation in the global promoter analysis (Extended Data Fig. 6b). This is indicative of local methylation alterations as the axolotl ages. In support of this, the methylation distribution analysis of highly methylated PRC2 targets suggests occurrences at 9.84 years old that could be interpreted as local hypermethylation (Fig. 5e, left panel). Overall, the consistency in correlation values and the methylation distribution across ages indicate that the methylation profiles, both at PRC2 target genes as well as across all promoter regions, remain stable.

Next, we aimed to shed light on the age-related methylation changes by performing EWAS of age targeting all promoter regions and segregating PRC2 target genes (Fig. 5g). Notably, the most significant age-related methylation changes were not linked to PRC2 targets, as indicated by the distribution of Z-scores (Fig. 5g). Moreover, the median Z-scores for both the overall promoter analysis and PRC2 targets-only analysis were close to zero, suggesting that methylation profiles of only a small proportion of genes are associated with age. Restricting the analysis to post-maturation samples results in higher positive and negative Z-scores. We hypothesized that by eliminating the youngest sample, we created an environment where slight changes in methylation could generate higher Z-scores (Fig. 5g, right panel). This suggests that most methylation changes occur during maturation. The variability introduced by the youngest sample could mask smaller, yet significant, changes in older samples, potentially skewing the profile for detecting Z-scores. However, this could be also due to the limited number of samples, which may reduce the statistical power of the results. Taken together, these findings are noteworthy, as they highlight axolotls may not exhibit classical epigenetic ageing traits seen in other species.

To challenge our clock findings, we analysed nanopore methylation profiles further. This time we selected promoters specifically related to CpGs from the methylation array analysis which were used to build the axolotl epigenetic clock (hereafter referred to as ‘clock promoters’). In the combined correlation analysis of all and top 100 age-associated clock genes along with hierarchical clustering, the strongest correlation was observed between the 3.55- and 9.84-year-old samples, indicating that their methylation profiles are the most similar (Extended Data Fig. 6d-f). This further supports the hypothesis that, following the early life period, the axolotl methylome undergoes fewer epigenetic modifications, in contrast to the dynamic changes occurring during the development-maturation phase. Independent analysis of all genes that are differentially methylated during maturation (in the youngest samples) showed they remain stable in the oldest samples (Extended Data Fig. 6c). Altogether, these findings corroborate the observations made with the Infinium arrays and the global methylation analyses, independently suggesting that axolotls exhibit epigenetic stability after maturation and thereby providing a molecular explanation for their negligible senescence.

## Discussion

This study presents the first epigenetic clocks for the axolotl, a species of outstanding biological relevance due to its regenerative abilities and negligible senescence features. Importantly, we uncover a hitherto unprecedented phenomenon, namely that the clocks are only able to capture age-related changes until 4 years of age (constituting the first third of the average axolotl lifespan), while no predictors can be built using samples spanning the entire species lifespan or for animals above 4 years of age. This is suggestive of an epigenetic switch taking place in axolotls at the end of the early life period which supports the stabilization of age-related methylation states, leading to the acquisition of negligible senescence at the epigenetic level.

This is all the more remarkable in light of several lines of evidence suggesting that the epigenetic ageing observed in axolotls during the early life period is evolutionarily conserved. First, we are able to build dual species clocks between early life axolotls and either frogs or humans based on a single mathematical formula, indicating the existence of shared methylation trends at highly conserved CpGs which are able to inform on epigenetic ageing across species. This also highlights the utility of our single and dual clocks as resources for ageing studies in this remarkable species. Second, our EWAS of age shows that top age-correlated CpGs during the axolotl early period include developmental and transcriptional regulators, including homeobox transcription factors, a major feature of mammalian and amphibian ageing^33,36,42,43^ which is even observed in evolutionarily distant zebrafish^31^. This is further confirmed by the two-way analysis comparing axolotl against human or frog age-related CpGs, and supported by our GREAT and GO enrichment analyses, which again highlight shared ageing trends between axolotls in their early life period and other species.

The generation of the clocks was enabled by the use of methylation arrays that profile CpGs which are highly conserved across mammals. We selected this platform as it was previously used to build accurate age predictors for frogs^33,34^, the phylogenetically closest group to salamanders. Yet, this approach is not without limitations, as the analysis is circumscribed to a subset of 5386 CpGs which map to the axolotl genome. This raises the possibility that our inability to build clocks for axolotls beyond the early life period is due to the exclusion of CpGs that may change throughout lifespan. However, we note that recently presented frog clocks were built with a smaller number of CpGs that align to the respective genomes providing evidence that the proposed model is applicable to track age across lifespan in amphibians. Moreover, this is further mitigated by the results stemming from our unbiased ONT genome-wide profiling of methyl cytosines within promoter regions, which indicates that a) 3.55- and 9.84 years-old individuals exhibit the highest similarity when considering differentially methylated promoters in the maturation period, b) there is a remarkable stability of the methylome at global level, including for key age-related sites such as those related to PRC2 targets (22% of the aligned CpGs correspond to bivalent promoters and ‘Repressed by Polycomb’ regions) and c) for all clock promoters, the closest methylation profiles are shared between the 3.55- and 9.84 years-old individuals, supporting the notion of methylome stabilization after the early life period. These considerations, together with the numerous evolutionarily conserved features of epigenetic ageing in axolotls up to 4 years of age which are not detectable afterwards, support the existence of a true divergence point towards methylome stability between early and late life individuals.

The biological processes that underlie this switch are as yet unclear. The divergence point does not coincide with sexual maturation (attained between 10-12 months of age^21^), nor with reported growth patterns^22^. Another possibility is that axolotls are neotenic organisms, meaning they retain juvenile traits while becoming sexually mature^21^, which may impact on clock behaviour. However, epigenetic clocks have been developed for naked mole rats (up to 26 years), mammals which also exhibit neoteny^43^. Whether the observed epigenetic stabilisation is associated with a lack of completion of the developmental programme in neoteny, or instead constitutes a trait of all salamander species, which exhibit negligible senescence, remains a critical outstanding question.

At the molecular level, this stability is manifested by a remarkably stable methylome which, in contrast to other species^36,37,54–56^, does not exhibit prominent age-related shifts towards hypo or hypermethylation. Further, we observed very weak methylation gains at PRC2 target sites in various tissues throughout axolotl lifespan. As methylation increases at such sites and a shifting DNA methylome constitute features of ageing across species^36,37,54,55^, our observations support the existence of true negligible senescence at the epigenetic level in the axolotl. We have recently shown that the rate of change in PRC2 target sites exhibits a strong negative correlation with maximum mammalian lifespan^65^. Thus, the very slow age-related gain in methylation at PRC2 sites suggests a long lifespan for the axolotl. Further studies into the role and features of epigenetic modifiers in this species could offer insights into the mechanisms underlying this striking stability.

The extent to which the remarkable regenerative abilities found in salamanders contribute to negligible senescence remains unknown. The successful construction of age predictors for the early life period will enable explorations into how axolotl regeneration impacts on epigenetic age. As such, the axolotl DNAm clocks presented herewith constitute a critical resource towards endeavours addressing this important question.

One last consideration is that axolotls may still be undergoing maturation until 4 years of age, with subsequent stabilisation of mature features and negligible senescence. In light of theories proposing the dysregulation of developmental circuits as a driver of ageing^66^, it is possible that the methylome stabilisation seen in mature axolotls is linked to an active regulation of developmental pathways (e.g., PRC2-related), a hypothesis which should be tested by future research. The latter would benefit from integrating DNAm profiles with transcriptomic and epigenomic datasets, which would provide a functional dimension to the analysis.

### Limitations of the study

In this study, we built accurate pan- and single tissue epigenetic clocks for early life axolotl using several organs. Although we used a large number of samples in the pan-tissue clock, single tissue limb and tail clocks are built using more restricted numbers, which may result in higher measurement errors. Larger sample sizes could be used to enhance the predictive power of such clocks.

Beyond array-based analysis, we also conducted genome-wide ONT sequencing to investigate methylation across promoter regions. Due to the features of this technology, we were limited to profiling a limited number of samples. Despite robust epigenetic assessment and high coverage, the small sample pool may affect the interpretation of results.

Loss of DNA methylation at super-enhancers is a well-characterised occurrence during mammalian ageing^67^. Future work should address whether this takes place in the axolotl.

While bulk analysis provides a representation of global tissue profile, this approach does not inform on population heterogeneity and individual cell susceptibility to ageing. Single cell studies would help dissecting smaller-scale contributions and determine the cell type drivers of the tissue clocks.

## Methods

### Ethics and Animal Husbandry

All animal procedures were conducted in compliance with the current German Animal Welfare Act and legislation from the state of Saxony, under the licence TVT 5/2021 issued by the animal welfare authorities.

Axolotls (*Ambystoma mexicanum*) were bred and maintained at the CRTD facility (Dresden, Germany). Axolotls were housed in individual aquaria at ∼18–20 °C and a 12h/12h light/dark cycle. Tissue samples from axolotls of the wild type strain, d/d (leucistic) lines and transgenic reporter models not expected to influence animal health were used in this study.

### Tissue Collection

For tissue collection, axolotls were sacrificed by 0.1 % benzocaine (Sigma) overdose. Solid tissues (forelimb, tail, liver and spleen) were collected by dissection and snap-frozen in liquid nitrogen before storage at −80 °C. Fresh blood was obtained by cutting external gills, and skin mucus was collected using 10 µl inoculation loops (Sarstedt) by applying 4-5 longitudinal movements along the axolotls’ tail and up to the back. Blood and mucus samples were collected in the digestion buffer and processed immediately.

### Cell Culture

Axolotl AL1 cells^68^ were grown on 0.75 % gelatin-coated plastic dishes in MEM (Gibco) supplemented with 10 % heat-inactivated foetal calf serum (FCS, Gibco), 25 % H2O, 2 nM L-Glutamine (Gibco), 10 μg/ml insulin (Sigma) and 100 U/ml penicillin/streptomycin (Gibco) in a humidified atmosphere of 2.5 % CO2 at 25 °C. Cells were sub-cultured using standard protocols as previously described^19^.

### Decitabine Treatment

For *in vitro* analysis, non-toxic concentrations of decitabine (5-aza-2’-deoxycytidine, DAC, Selleck Chemicals, Cat# S1200) were determined by alamarBlue viability assay (BioRad) using a Promega GloMax platereader, after 4-days treatment. Subsequently, AL1 cells were treated with 0.05, 0.1 or 1 μM decitabine or DMSO vehicle control and cultured in triplicates (n = 2 in the cells batch treated with 0.05 μM decitabine). Drug-supplemented media was administered for 7 days. Cell pellets were collected by centrifugation, fast-frozen in liquid nitrogen and stored at −80 °C until processing.

### gDNA Isolation

Genomic DNA extraction was performed using Quick-DNA™ Miniprep Plus Kit (Zymo Research, Cat# D4069) following manufacturer’s instructions with several adjustments. Solid and liquid tissue samples were digested overnight at 55 °C. Thawed axolotl AL-1 cells were resuspended in 100 μl of DNA Elution Buffer prior to digestion and were digested for 3-5 hours. DNA was purified and concentrated using DNA Clean & Concentrator-5 Kit (Zymo Research, Cat# D4014) following the manufacturer’s protocol.

### Relative Global Methylation Assessment

5-mC DNA ELISA Kit (Zymo Research, Cat# D5326) was used to detect the global 5-methylcytosine (5-mC) in DNA, following the manufacturer’s protocol. All samples were analysed in technical duplicates. For demethylation treatment and age-related methylation analysis, 100 ng DNA per sample was utilised. The absorbance was measured with a Promega GloMax plate reader or FlexStation 3 microplate reader. The obtained absorbance values were utilised to estimate changes in relative global methylation levels. Relative global methylation values were calculated by normalising them with the methylation levels observed in the control samples for AL-1 cells drug treatment experiments or in tissues from the youngest animals (for the estimation of global methylation across ageing).

For the statistical analysis, we employed one-way ANOVA with Tukey multiple comparisons test of means.

### DNA methylation arrays

All DNA methylation data were generated using the mammalian Infinium array “HorvathMammalMethylChip40”^44^. By design, the mammalian methylation array facilitates epigenetic studies across mammalian species (including humans) due to its very high coverage (over thousand-fold) of highly-conserved CpGs in mammals. A subset of cytosines on the mammalian array also apply to more distant species including amphibians^44^.

The Infinium method is based on sodium bisulfite conversion of DNA and microarray-based genotyping of CpG sites using Infinium bead technology with single-base resolution. The advantage of the microarray platform is that it is user-friendly, it can be multiplexed, and it exhibits good agreement with other platforms’ DNA methylation measures. Specifically, the Infinium beads bear a 23-base oligo address to locate them on the BeadChip, and a 50-base probe. Probe sequences are complementary to specific 50 base regions of bisulfite-converted genomic DNA. The 3′ end of the probe harbours the methylated CpG site to be monitored. After the probe is hybridized to bisulfite-treated test DNA, a single-base extension adds a fluorescently labelled ddNTP to the 3′ CpG site. This lets the C to T change caused by bisulfite conversion to be “genotyped.” The fluorescent signal is then measured and processed.

CpG annotation was performed as described^27,44^. Out of all CpGs on the mammalian array, 5,386 CpGs map to the axolotl genome according to genome assembly AmexG_v6.0-DD from UCSC. Genome coordinate information can be downloaded from our GitHub page (https://github.com/shorvath/MammalianMethylationConsortium) and the supplementary information in^44^. The chip manifest file can be found at Gene Expression Omnibus (GEO) at NCBI as platform GPL28271. The SeSaMe normalization method was used to define beta values for each probe^69^.

### Penalised regression models

Details on the clocks (CpGs, genome coordinates) and R software code are provided in Supplement. Penalized regression models were created with glmnet^70^. We investigated models produced by “elastic net” regression (alpha=0.5). The optimal penalty parameters in all cases were determined automatically by using a 10-fold internal cross-validation (cv.glmnet) on the training set. By definition, the alpha value for the elastic net regression was set to 0.5 (midpoint between Ridge and Lasso type regression) and was not optimized for model performance.

We performed a cross-validation scheme for arriving at unbiased (or at least less biased) estimates of the accuracy of the different DNAm based age estimators. For validation of the clocks, we used leave-one-out LOO cross-validation (LOOCV) in which one sample was left out of the regression, then predicted the age for the remaining samples and iterated this process over all samples.

A critical step is the transformation of chronological age (the dependent variable). We used a simple modified log transformation for all of the clocks for axolotls presented in this study (namely, ln(age + 0.5)). For the details of how the age transformation is used in the definition of the epigenetic clocks, see Supplement.

The decision to try an age-restricted clock came from an observation that, when building all-age clocks, that prediction improved as more extreme log-type concave age transformations were used. We noticed that LOO correlation across all 4 tissue-defined clocks improved to different degrees, depending on the specific age cut-off chosen. In order to select the final age cut-off of 4.0 years, we examined the LOO correlation for all types of standard age transformations (from square root to log linear), with different age cut-offs ranging from 1.0 to 7.0 years, in increments of 0.5 years. Age cut-offs beyond this range were ruled out due to a trend of extremely poor prediction. The final age cut-off was chosen as the best choice when weighing all 4 clocks together.

It is important to note that the set of CpGs used to train the Limb-specific clocks is not the same set of CpGs that map to the axolotl genome according to genome assemblies. Rather, for the Limb-only clocks, the CpGs presented to the regression model were those that were considered detectable, based on mean methylation values (Supplement). The reason that this narrower filter was applied only to the Limb-specific clocks is because it improved the prediction and performance of the Early-Late life Limb-specific clock during cross-validation analysis, where as it negatively affected prediction among the other three Early-Late life clocks.

### EWAS of age

We conducted an epigenome-wide association study (EWAS) of age on 5,386 CpGs mapped to the axolotl genome. In our EWAS analysis, we restricted the axolotl samples to those with a chronological age of ≤4 years. The analysis was performed across all tissue types (n=97), limbs and tails only (n=46), and specific tissue types: blood (n=13), limb (n=19), liver (n=10), skin (n=17), spleen (n=11), and tail (n=27). Association analysis was conducted using the standardScreeningNumericTrait function under R WGCNA. To estimate the association between chronological age and methylation levels, we employed a robust correlation test (biweight midcorrelation, bicor), which is less sensitive to outlier data points^71^. A suggestive significance threshold was set at P<1.0E-04.

### Enrichment analysis for annotating EWAS of age

Similar to our previous mammalian aging study^36^, we annotated age-related CpGs based on chromatin state analysis to identify enriched states in the axolotl aging process and conducted GREAT analysis to identify biological pathways and functional gene sets.

### Chromatin state analysis

For chromatin state annotation, we employed a recently published universal ChromHMM chromatin state annotation of the human genome^72^. The universal ChromHMM model covers a total of 100 distinct states, categorized into 16 major groups in the human genome. Each axolotl-mapped CpG (n=5,386) was assigned a chromatin state based on hg19, and we only analyzed states with at least 20 CpGs. This yielded a total of 64 states across 13 groups, including quiescent, heterochromatin (HET), polycomb repressed, acetylations, weak enhancers, enhancers, transcribed and enhancer, weak transcription, transcription, exon, weak promoters, promoters, and transcription start site (TSS), available for analysis. For annotation, we used the top 150 CpGs with positive and negative age correlation from our EWAS of age, respectively.

### Overlap with Polycomb Repressive Complex Binding Regions

Polycomb repressive complex (PRC) annotations were defined as in our previous study^36^. Briefly, annotations were based on the binding of at least two transcription factor members of polycomb repressor complexes PRC1 (subgroups RING1, RNF2, BMI1) or PRC2 (subgroups EED, SUZ12, EZH2) using 49 available ChIP-seq datasets from ENCODE^73^.

In the current study, we applied the same PRC2 annotations to analyze the overlap with age-related CpGs. Of the 5,386 axolotl-mapped CpGs, 39 CpGs are annotated for PRC1 and 634 CpGs are annotated for PRC2.

In both chromatin state and PRC annotation analysis, we conducted a one-sided hypergeometric analysis to examine enrichment (Odds ratios [OR] >1) and depletion (OR <1) patterns, focusing on the top 150 CpGs that increased with age and the top 150 CpGs that decreased with age from our EWAS of age.

PRC2 target sites selection is based on human cells annotation identified by ChIP-seq. Human datasets are used due to the high level of evolutionary conservation of Polycomb complexes and their targets as shown in mammalian^36^ and amphibian species^33^. In the axolotl datasets, genomic regions predicted to be PRC2 binding sites largely display hypomethylated states, which aligns with our expectations. This forms the basis for using the current annotation in axolotls.

### GREAT analysis

We conducted GREAT enrichment analysis^51^ on the top 150 positively age-related and the top 150 negatively age-related CpGs identified from our EWAS of age. The GREAT framework implements foreground/background hypergeometric tests that are not confounded by the number of CpGs within a gene or gene size. For the analysis, we used all 5,386 axolotl-mapped CpGs as the background and the genomic regions of the top 150 CpGs as the foreground. The enrichment analysis parameters were set as follows: human genome assembly hg19, proximal regions defined as 5.0 kb upstream and 1.0 kb downstream, and distal regions extending up to 50 kb, consistent with the methodology employed in our previous mammalian aging study^36^. The GREAT analysis was executed using the rGREAT package in R. We report gene sets with FDR < 0.05, nominal hypergeometric P-values < 0.001, and a minimum of 3 overlapping genes. FDR calculations were performed for each ontology, including GO, MSigDB (with upstream regulator gene sets), PANTHER, KEGG pathway, disease ontology, gene ontology, human phenotype, and mouse phenotype.

### EnrichR analysis

Gene set enrichment analyses of the top 196 age-related CpGs (filtered with pvalueStudent <0.01) in the pan-tissue dataset was also performed using enrichment annotation tool EnrichR (EnrichR package in R^74^).

The analysis was performed using default settings, with human genes selected as background. The following gene set libraries were selected: GO – GO Biological Process 2023 ontology, R – Reactome 2022 ontology. The EnrichR tool was employed to calculate p-value with adjustments for multiple comparisons using the Benjamini-Hochberg method. Enrichment terms were filtered based on the adjusted p-value < 5e-2. We calculated the richFactor values, which is the ratio of the enriched genes in the term to the total number of genes assigned to the term. We report the negative log transformed adjusted p-value (logP = - log10(Adjusted.P.value)).

### Nanopore-based methylation analysis

For the analysis, we used limb tissue from one animal per age point (0.73, 1.1, 3.55 and 9.84 years old). DNA was isolated using Quick-DNA™ Miniprep Plus Kit (Zymo Research, Cat# D4069) as described above. Input into library construction was 4ug gDNA, fragmented to 6-7kb with a gTUBE (Covaris). Libraries were prepared using ligation sequencing kit V14 SQK-LSK114 (Oxford Nanopore Technologies) and loaded onto 3-4 R10.4.1 PromethION FLO-PRO114M flow cells per sample. Each flow cell was flushed using wash kit EXP-WSH004 and reloaded after approximately 6 hours of sequencing due to high rates of pore blocking. Sequencing was performed using adaptive sampling to enrich for promoter sequences in the genome. A bed file was created selecting 87,931 regions of 10kbp each, some of these regions overlapped leading to a total selected size of 861,267,254 bps. Reads were basecalled using dorado (v0.3.3) and 5mC at CpG contexts were called using <u>dna_r10.4.1_e8.2_400bps_sup@v4.2.0_5mCG_5hmCG</u>. Despite the adaptive sampling, individual CpG coverage was still low at a mean of 13x at the targeted regions, therefore we combined all CpGs within the promoter region of a gene (defined as +/-5kbp) to produce a single methylation value for each promoter.

### Methylation distribution for nanopore data

The methylation dataset generated by nanopore sequencing was analysed using R software. Prior analysis, AMEX gene regions with methylation = 0 in all age points were filtered out. High-methylation promoter regions were filtered based on a methylation value > 0.7 in all ages, while a value of <0.3 was used for low-methylation ones. For methylation distribution plots we report median.

To identify PRC2 target genes among promoter regions we applied the same PRC2 annotations from the MSigDB database as described above^51,60,75^.

For the statistical analysis, we employed Kruskal-Wallis test with adjustments for multiple comparisons using the Bonferroni method.

### Correlation analysis for nanopore data

Correlation analysis was performed using the R function “cor” (method = “pearson”) and correlation matrices were visualised using the R function “corrplot” (type = “lower”, method = “square”, order=”hclust”) from the R package “corrplot”.

### EWAS of age for nanopore data

We conducted EWAS of age on average methylation values across all promoters. Association analysis was conducted using the standardScreeningNumericTrait function under R WGCNA^71^.

### Differential methylation analysis for nanopore data

Differentially methylated promoter regions in the youngest samples were calculated by subtracting methylation values between samples (distance [gene 1] = age0.73 [gene 1] – age1.1 [gene 1]. If a calculated distance was > 0.2 or < −0.2, the promoter region was considered as differentially methylated between 0.73- and 1.1-years old samples. Differentially methylated promoter regions were then filtered out and used for the correlation analysis between all ages.

### Heatmap clustering for nanopore data

Clock promoter regions were filtered based on CpGs from the methylation array analysis which align to axolotl genome. The top 100 clock promoter regions were selected based on the top 100 age-associated CpGs from the EWAS of age (pan-tissue, methylation array dataset). Heatmap was built using the R function “pheatmap” (cutree_rows = 6, cluster_rows = T, cluster_cols = T, clustering_distance_cols = “euclidean”) from the R package “pheatmap”.

## Data availability

The human and frog methylation datasets used herewith were previously published and publicly available^33,46^. The mammalian methylation array is accessible via the non-profit Epigenetic Clock Development Foundation (https://clockfoundation.org).

## Code availability

The R software code used for clock development is provided in the Supplement.

## Acknowledgements

We thank all members of the Yun lab and Ken Raj for comments, Anna Czarkwiani, Nina Loetzsch and Mark Seiferth for tissue collection assistance, Anja Wagner and Beate Gruhl for axolotl husbandry. This work was supported by a DAAD and a Saxon Scholarship to Y.H., an Alexander von Humboldt postdoctoral fellowship to H.E.W., and DFG (22137416, 450807335, and 497658823) grants, CRTD and ALTOS funds to M.H.Y. S.H. and the Altos team were funded by Altos Labs.

## Author contributions

Y.H., H.E.W., M.L., N.P., and M.H.Y generated DNA samples, designed and performed experiments. J.A.Z., S.H., Y.H., A.H., A.L. and R.L. performed statistical analyses. All authors analysed and interpreted data. M.H.Y. and Y.H. drafted the manuscript. All authors edited the manuscript. M.H.Y and S.H. conceived and supervised the study.

## Additional information

**Extended data Figures 1 - 6**

## Supplementary information

**Supplementary Table 1_Description of Axolotl data by tissue type**

**Supplementary Table 2_Clock coefficients and genome annotation**

**Supplementary Table 3_CpGs identified in Axolotl EWAS of age**

## Supplementary material

**Correspondence and requests for materials should be addressed to**

---

**Extended Data Fig 1.**
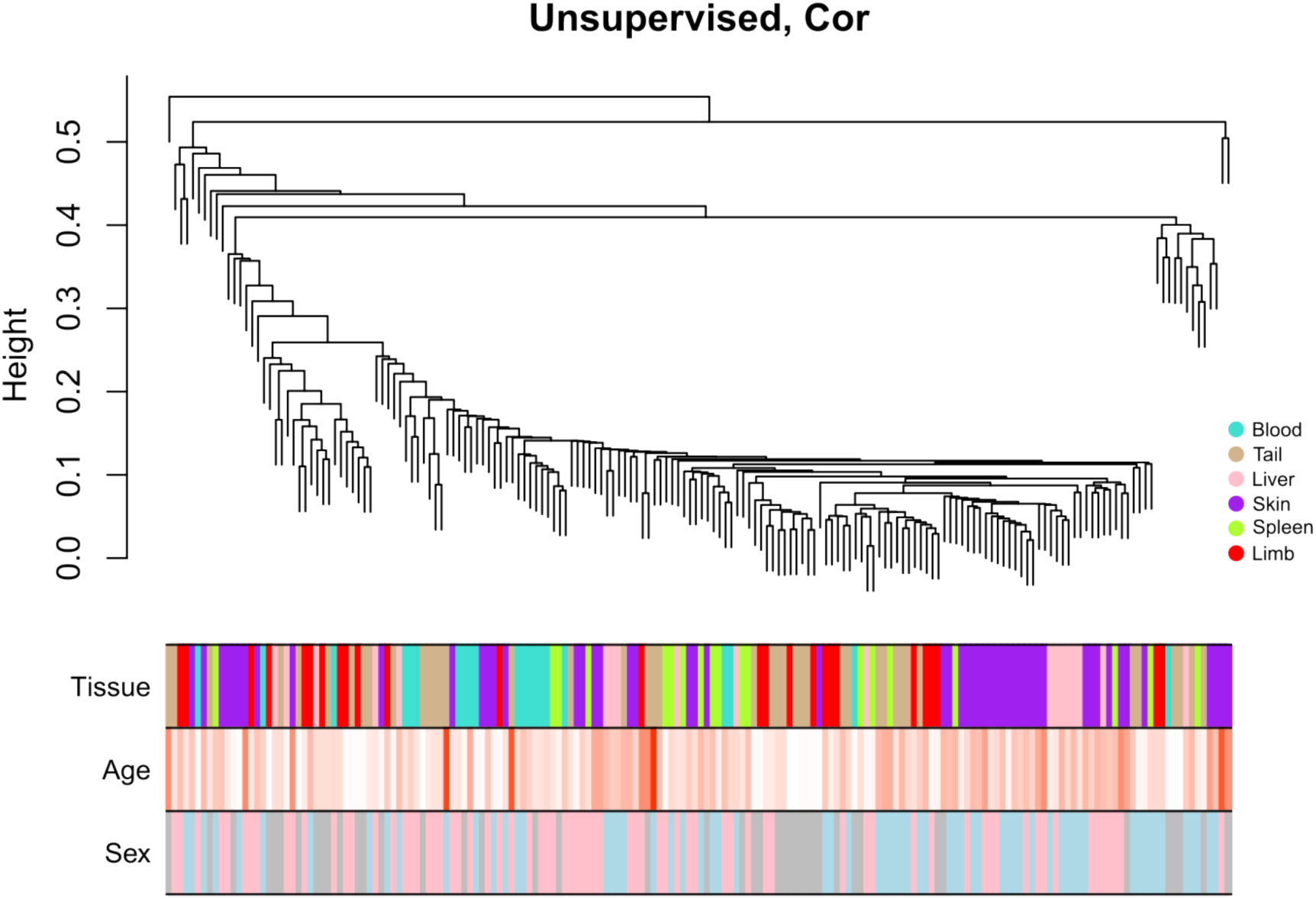
Unsupervised hierarchical clustering of *A. mexicanum* tissues. The clustering height (y-axis) can be interpreted as distance based on pairwise correlation coefficients (1-cor based on detectable CpGs). Colour bands underneath indicate clustering branch corresponding to a height cut-off of 0.6 on the y-axis. Total sample size: n = 180 samples. Tissue type: tan = tail (n = 42), red = limb (n = 26), purple = skin (n = 53), turquoise = blood (n = 21), green yellow = spleen (n = 19), pink = liver (n = 19). Age is coloured as follows: red corresponds to old age (oldest = 21 years) and white corresponds to the young age (youngest = 0.08 years). Sex is coloured as follows: pink - female, blue - male, grey - unknown (animals were too young to determine their sex). Samples don’t show clear separation by age or tissue.

**Extended Data Fig 2.**
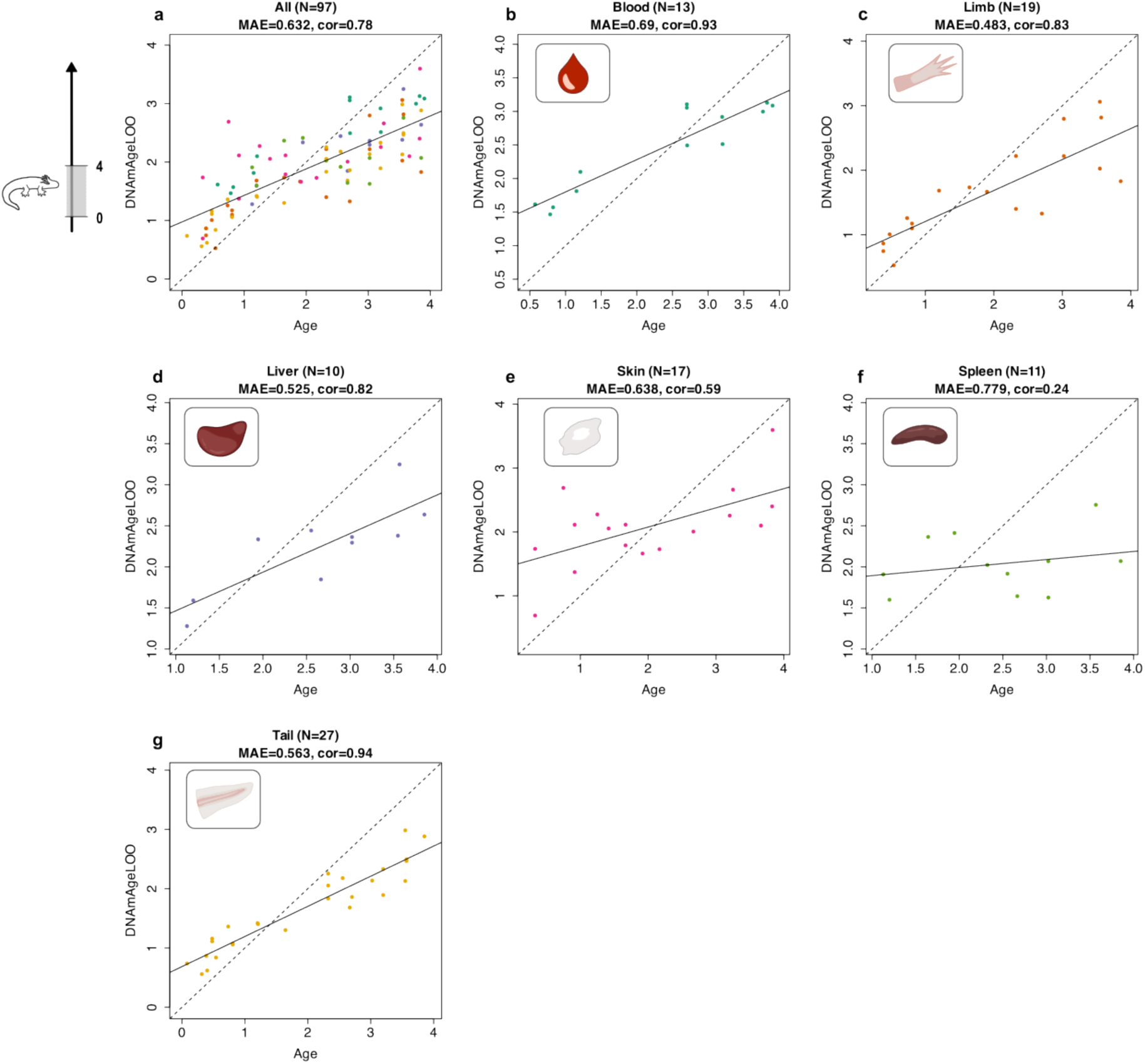
Early life axolotl pan-tissue epigenetic clocks, plotted by tissue of origin. Cross-validation study of epigenetic clocks for *Ambystoma mexicanum*. Leave-one-sample-out (LOO) estimate of DNA methylation age (y-axis, in units of years) versus chronological age (x-axis, units in years) for *A. mexicanum* samples across selected range of age. Dots are coloured by tissue type. **a**-**g**, Pan-tissue epigenetic clock for the early life cohort (<4 years), represented by tissue subset: **a**, pan-tissue, **b**, blood, **c**, limb, **d**, liver, **e**, skin, **f**, spleen, **g**, tail. Each panel reports the total sample size (N), median absolute error (MAE), in years, Pearson correlation coefficient (cor).

**Extended Data Fig. 3.**
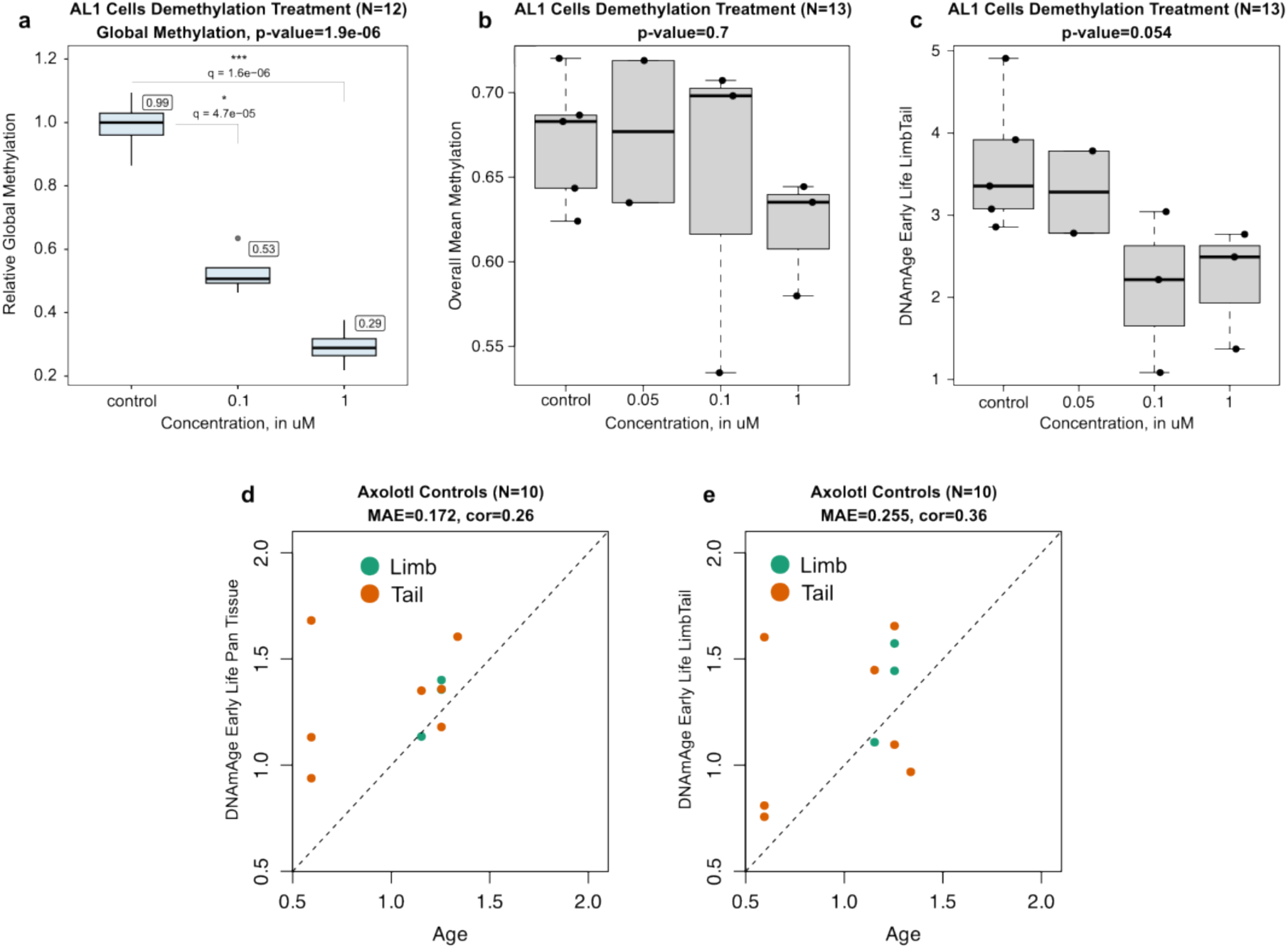
Axolotl epigenetic clocks reflect changes in methylation status upon interventions and predict age of samples from independent datasets. **a-c**, Assessment of Infinium array’s sensitivity to methylation changes in axolotl cells induced by exposure to hypomethylating agent decitabine (DAC). In box plots, the centre line denotes median. Means are presented in white boxes. Each panel reports the total sample size (N) and p-value. **a**, Relative global methylation changes measured by 5-mC DNA ELISA-based method in axolotl AL-1 cells exposed to the indicated concentrations of DAC. Statistical significance was evaluated by one-way ANOVA with adjustments for comparisons between treated cells groups and control using the Tukey method, p-value and q-values are indicated. **b**, Total methylation values extracted from Infinium methylation arrays for samples derived from AL-1 cells treated as in **a**. **c**, DNAm age changes in AL-1 cells for the indicated conditions. Statistical significance in **b** and **c** was evaluated by Kruskal-Wallis test. **d**-**e**, Leave-one-sample-out (LOO) estimate of DNA methylation age (y-axis, in units of years) versus chronological age (x-axis, units in years) for *A. mexicanum* samples corresponding to the early life age cohort (<4 years). Dots are coloured by tissue type. **d**, pan-tissue epigenetic clock, **e**, Limb-tail epigenetic clock. The panel reports the total sample size (N), median absolute error (MAE), in years, Pearson correlation coefficient (cor).

**Extended Data Fig 4.**
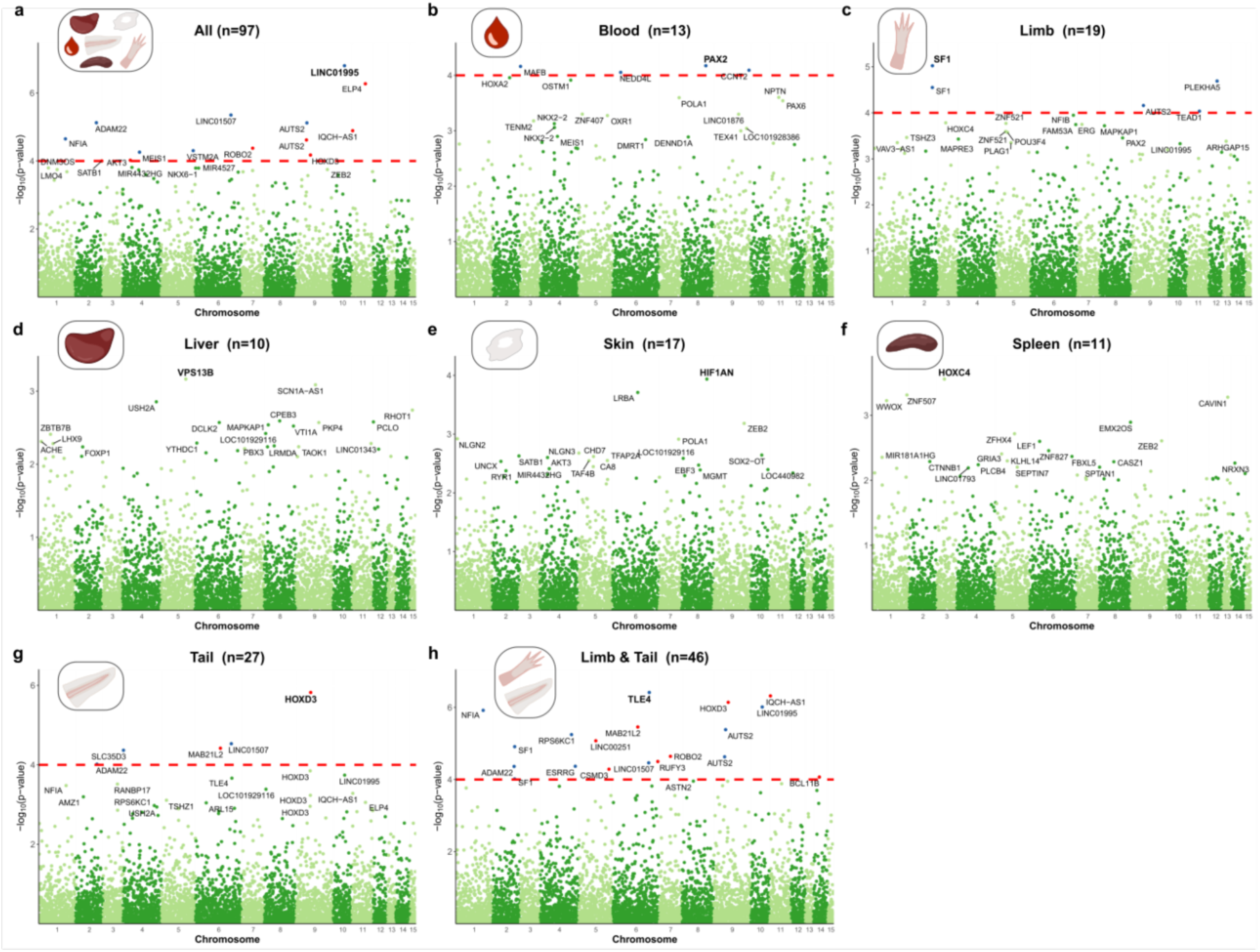
Meta-analysis of age-related methylation change during axolotl early life. **a**-**h**, Manhattan plots of the epigenome-wide association (EWAS) of chronological age in *A. mexicanum* tissues during early life (<4 years): **a**, pan-tissue, **b**, blood, **c**, limb, **d**, liver, **e**, Skin, **f**, Spleen, **g**, tail, **h**, limb-tail. The red dash lines indicate suggestive levels of significance at P < 1.0E-04. CpGs in red and blue reflect positive and negative correlation with age, respectively. The y-axis displays -log base 10 transformed P-value and the x-axis displays chromosome number based on the *A. mexicanum* genome (v. 6.0-DD). Labels are provided for the top 20 hypermethylated/hypomethylated CpGs according to the product of Z scores in x- and y-axis. Each panel reports the total sample size (n).

**Extended Data Fig. 5.**
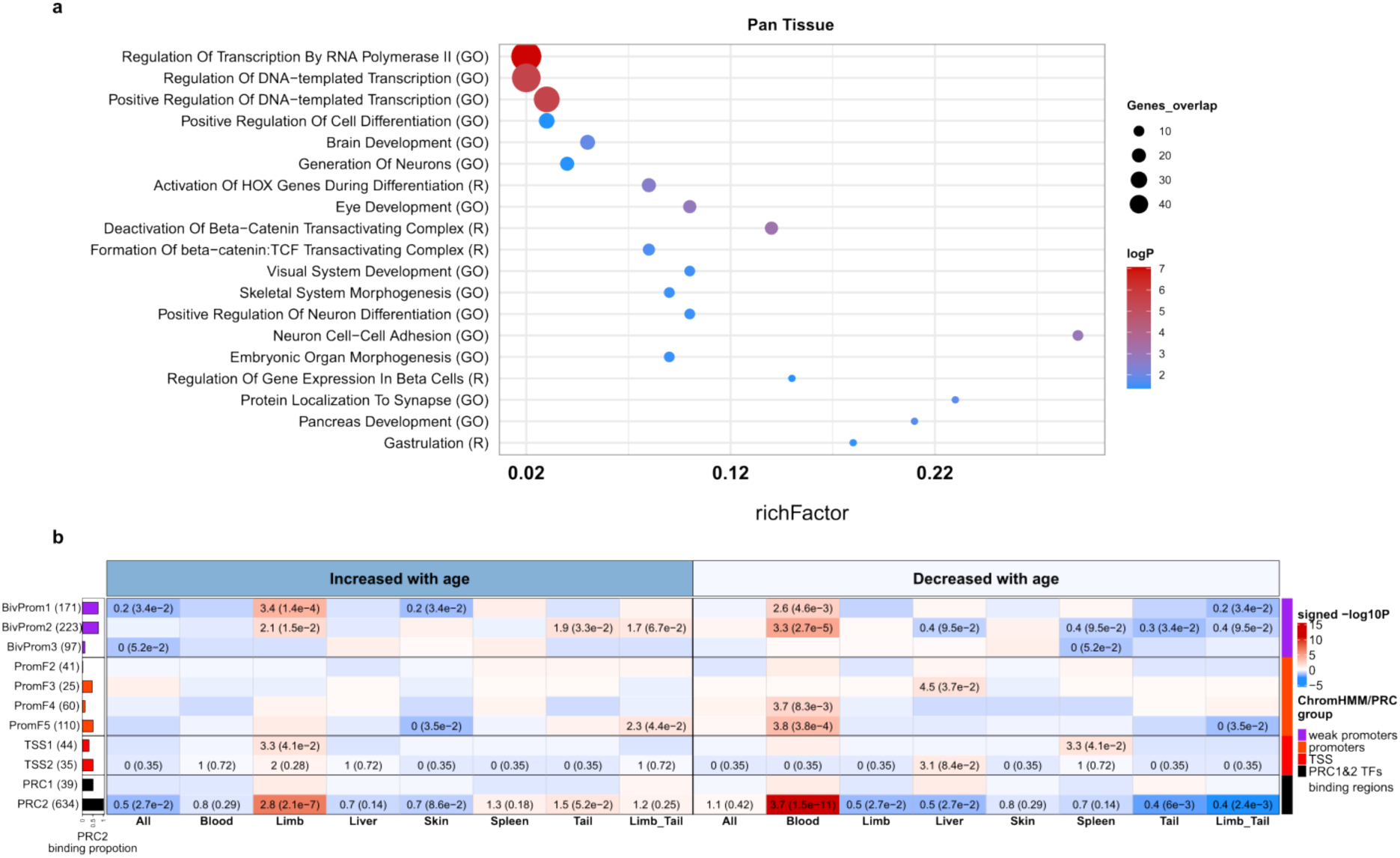
Functional enrichment of age-associated CpGs highlights developmental gene signatures during early life changes in axolotl tissues. **a**, Functional enrichment results for top age-related CpGs from pan-tissue EWAS. The y-axis denotes the name of a functional gene set/biological pathway and the ontology. The x-axis denotes the richFactor showing the degree of enrichment in respective ontology terms. The dot colour gradient is based on -log10 (adjusted P-value). **b**, Chromatin state analysis of age-related CpGs from EWAS in 1) pan-tissue, 2) blood, 3) limb, 4) liver, 5) skin, 6) spleen, 7) tail, 8) limb-tail, respectively. The y-axis denotes chromatin states and Polycomb repressive complex 1 and 2 (PRC1, PRC2) enrichment. The analysis is based on Polycomb target sites ChipSeq datasets in ENCODE^73^ and universal chromatin states annotation^72^. The heatmap colour gradient is based on -log10 (unadjusted hypergeometric P-value) multiplied by the sign of OR > 1. Red and blue colours denote OR > 1 and OR < 1, respectively. Abbreviations: ChromHmm – software for learning chromatin state, BivProm1-3-bivalent promoter state associated with promoter marks and H3K27me3, PromF2-5 - flanking promoter state, TSS1-2 - transcription start sites, TF - transcription factors, GO – GO Biological Process 2023 ontology, R – Reactome 2022 ontology.

**Extended Data Fig. 6.**
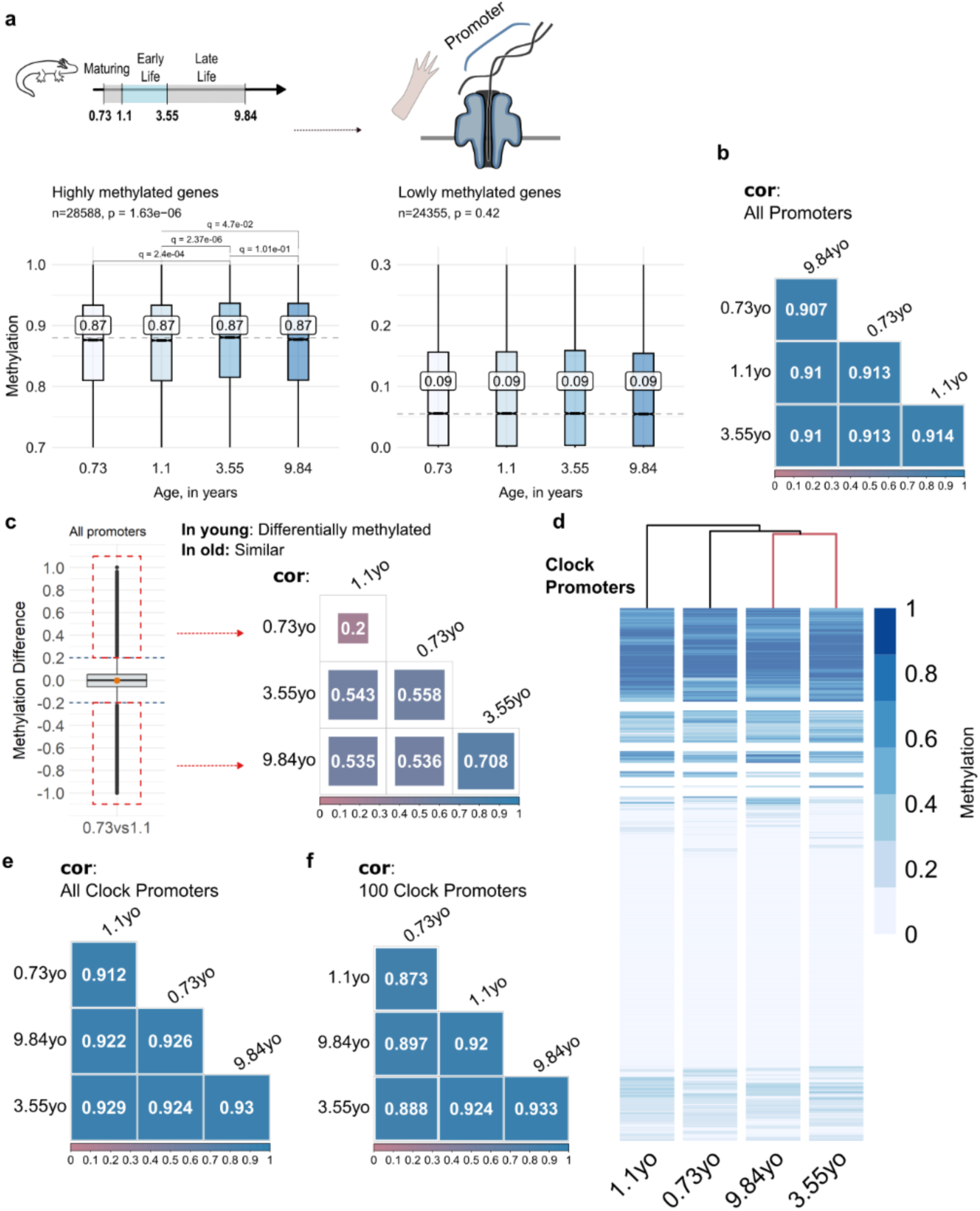
Nanopore-mediated adaptive sampling reveals stability in the epigenetic landscape across axolotl ageing. **a**, Top: schematic representation of the experiment set up. Promoter-targeted nanopore sequencing of axolotl limb samples with methylation calling. The age of the limb samples is indicated in years. Bottom: Box plots of the methylation distribution in detected promoter regions across axolotl limb samples selecting highly (β-value > 0.7) and lowly methylated (β-value < 0.3) genes. Statistical significance was evaluated by Kruskal-Wallis test with adjustments for multiple comparisons using the Bonferroni method, p-values and q-values are indicated. Box plots center line reports median. Means are presented in white boxes. Panels report the gene regions number (n). **b**, Age pairwise Pearson correlation analysis of detected promoters. Colours indicate the degree of pairwise correlation. **c**, Differential methylation analysis between younger and older age groups. The box plot illustrates the distribution of differentially methylated genes in younger age group – 0.73yo and 1.1yo. The y-axis represents the methylation difference, calculated by subtracting the methylation values of the youngest age group – 0.73yo - from the 1.1yo. The genes with a methylation difference greater than ±0.2 are included for the all-age groups pairwise Pearson correlation analysis. Colours indicate the degree of pairwise correlation. **d**-**f**, Methylation analysis targeting promoters based on CpGs that are present on the mammalian array which map to the *A. mexicanum* genome. These promoters are referred to as ‘clock promoters’. **d**, Heatmap with dendrograms of average methylation levels of targeted clock promoters depicting hierarchical clustering of age groups. **e**-**f**, Age pairwise Pearson correlation analysis of **e**, all clock genes, and, **f**, top 100 age-related clock genes based on EWAS of age from the methylation array analysis. Colours indicate the degree of pairwise correlation.

