## Supplementary Material for "Axolotl epigenetic clocks offer insights into the nature of negligible senescence"

Here we present details on the construction and software for our epigenetic clocks for Axolotl

A) All Early Life Axolotl (age <= 4.0 years) tissues
B) Early life Axolotl (age <= 4.0 years) tissues, restricted to Limb and Tail
C) Early life Axolotl (age <= 4.0 years) tissues, restricted to Limb only
D) Early life Axolotl (age <= 4.0 years) tissues, restricted to Tail only

### Technical details surrounding epigenetic clocks

#### Statistical methods used for building the clocks

Since a mean methylation level of 0.5 is usually associated with a non-detectable CpG (i.e., a CpG whose sequence does not map to the species genome), we sometimes restricted the analysis to cytosines whose mean methylation across all Axolotl tissues was outside of [0.47,0.53], after the initial restriction to the 5,386 CpGs that map to the axolotl genome. These additional restrictions were only used when building the Limb-only clocks.

The epigenetic clocks were used by employing a single elastic net regression model analysis (R function glmnet). We use used Leave-one-out analysis (LOO) using a single lambda value. We chose the following parameters for the glmnet R function (Alpha: 0.5, CV Fold: 10, Lambda choice for Clock: 1 standard error above minimum CV-MSE).

#### Covariates and coefficient values of the axolotl clocks

The coefficient values of the clocks are specified in **Supplementary Table 2**.

1) The Early Life Axolotl clock (all tissues combined) is based on 49 CpGs whose coefficient values are specified in the column "Coef.AxolotlEarlyLife.Log2". Age transformation = ln( age + 0.5 ).
2) The Early Life Axolotl Limb-Tail clock (only samples from limb and tail tissues) is based on 49 CpGs whose coefficient values are specified in the column "Coef.AxolotlEarlyLifeLimbTail.Log2". Age transformation = ln( age + 0.5 ).
3) The Early Life Axolotl Limb clock (only samples from limb tissue) is based on 22 CpGs whose coefficient values are specified in the column "Coef.Axolotl EarlyLifeLimb.Log2". Age transformation = ln( age + 0.5 ).
4) The Early Life Axolotl Tail clock (only samples from tail tissue) is based on 22 CpGs whose coefficient values are specified in the column "Coef.AxolotlEarlyLifeTail.Log2". Age transformation = ln( age + 0.5 ).

#### General description of age transformation

An elastic net regression model (implemented in the glmnet R function) was used to regress a transformed version of age on the beta values in the training data. The glmnet function requires the user to specify two parameters (alpha and lambda). Since I used an elastic net predictor, alpha was set to 0.5. But the lambda value of was chosen by applying a 10-fold cross validation to the training data (via the R function cv.glmnet).

The elastic net regression results in a linear regression model whose coefficients b_0_, b_1_, . . . , relate to transformed age as follows
*F*(chronological age)=*b*_0_*+b*_1_*CpG*_1_*+ . . . +b*_p_*CpG*_p_+error

Note that the intercept term is denoted by b_0_. The coefficient values can be found in Supplementary Table 1. Based on the coefficient values from the regression model, DNAmAge is estimated as follows
*DNAm*Age=$F^{-1}$(*b*_0_*+b*_1_*CpG*_1_*+ . . . +b*_p_*CpG*_p_),

where $F^{-1}\left( y \right)$ denotes the mathematical inverse of the function F(.). Thus, the regression model can be used to predict to transformed age value by simply plugging the beta values of the selected CpGs into the formula.
